# Correlations in microbial abundance data reveal host-bacteria and bacteria-bacteria interactions jointly shaping the *C. elegans* microbiome

**DOI:** 10.1101/2024.06.13.598851

**Authors:** K. Michael Martini, Satya Spandana Boddu, Megan N. Taylor, Ilya Nemenman, Nic M. Vega

## Abstract

Compositional structure of host-associated microbiomes is potentially affected by interactions among the microbes and between the microbes and the host. To quantify the relative importance of these contributions to the microbiome composition and variation, here we analyze absolute abundance (count) data for a minimal eight-species native microbiome in the *Caenorhabditis elegans* intestine. We find that a simple neutral model only considering migration, birth, death, and competition for space among the bacteria can capture the means and variances of bacterial abundance, but not the experimental bacteria-bacteria covariances. We find that either bacteria-bacteria interactions or correlations among bacterial population dynamics parameters induced by the host can qualitatively recapitulate the observed correlations among bacterial taxa. However, neither model is uniquely or completely sufficient to explain the data. Further, we observe that different interactions are required to explain (co)variance data in microbiota associated with different host genotypes, suggesting different community dynamics associated with these host types. Finally, we find that many of these signals are obscured when data are converted to proportions from counts, consistent with a growing literature on the limitations of compositional data for inference of population dynamics. We end with discussing the limitations of Lotka-Volterra type assumptions for microbial community data analysis revealed by our results.

## Introduction

The drivers of community composition and individual variation in intestinal microbiota are not well understood (***Schlomann and Parthasarathy, 2019***). The composition is characteristic for hosts of a given type, with organisms from small minimal hosts (worms, fruit flies, etc.) to humans and other large mammals having host type-specific microbiota with a characteristic range of taxonomic and functional composition (***Skwara et al., 2023***; ***Burke et al., 2011***; ***Louca et al., 2016***). However, compositional variation across individual hosts is substantial (***Johnke et al., 2020***; ***Shreiner et al., 2015***; ***Burke et al., 2011***; ***Falony et al., 2016***), and mathematical models are essential to systematize this variability.

Unfortunately, the nature of microbiome data makes it difficult to accurately model the ecology of these systems. Indeed, generalized Lotka-Volterra (gLV) type models, where microbial taxa in a shared environment interact via a matrix *A* of pairwise terms during growth to saturation, have been used to approximate microbiome dynamics with some success (***Rakoff-Nahoum et al., 2021***; ***Joseph et al., 2020***). However, there are limitations to this approach when models are parameterized from data. First, microbiome data are in general obtained from community metagenomic sequencing and are therefore compositional; the caveats associated with fixed-count relative-abundance data are well described ***Joseph et al. (2020***); ***Gloor et al. (2017***); ***Tsilimigras and Fodor (2016***); ***Jian et al. (2020***). Unique identification of population dynamic parameters is often not possible from compositional data ***Remien et al. (2021***). There is considerable theory on the effects of data compositionality, but it is rarely possible to compare inference on real absolute-vs relative-abundance data, as absolute abundance is rarely measured in microbial community samples.

Further, to avoid over-fitting, it is generally necessary to assume that the parameters of these models are constant (time-independent) and are conserved across hosts of a given type (***Jones and Carlson, 2019***). This assumption is important—heterogeneity within data sets can readily obscure signals of interaction, resulting in incorrectly inferred networks (***Armitage and Jones, 2019***; ***Berry and Widder, 2014***)—but is difficult to test directly. Similarly, mathematical models often assume microbial interactions to be independent on the environment, such as the host background, while experiments suggest this assumption to be incorrect (***Bittleston et al., 2020***; ***Shetty et al., 2022***; ***Chamberlain et al., 2014***; ***Liu et al., 2017***; ***Deines et al., 2020***). The extent to which dependence of microbial interactions on time, environment, host genetic background, or even individual host properties affects the ability to use gLV mathematical models to make accurate inferences about the microbiome structure from data remains unknown.

Minimal microbiome systems provide an opportunity to address these questions directly. The nematode *C. elegans* has been recently developed as a model system for host-microbiome interactions (***Zhang et al., 2017***), for which diverse collections of culturable native microbiome isolates are available (***Dirksen et al., 2020, 2016***; ***Berg et al., 2016a***; ***Samuel et al., 2016***). In its natural environment, *C. elegans* contains characteristic communities of intestinal bacteria. These bacteria are obtained from the soil environment, but the composition of worm intestinal communities differs substantially from those found in the surrounding soil (***Dirksen et al., 2016***; ***Berg et al., 2016a***; ***Dirksen et al., 2020***; ***Johnke et al., 2020***). This suggests that the worm intestine provides a distinctive habitat and that, at a minimum, environmental filtering is important for determining the composition of these communities. Some of these microbes are transient mono-colonizers, unable to sustain populations without the continued arrival of new migrants, whereas others are able to proliferate independently in the intestine. However, persistence of individual strains increases when they colonize the intestine as part of a community (***Dirksen et al., 2020***). Differences in host genetics, in particular changes in immune function, are associated with changes in gut microbiome composition (***Berg et al., 2019***; ***Taylor and Vega, 2021***), suggesting that this small host actively controls its microbiome.

Even in this minimal system, where hosts from a shared genetic background are colonized from a shared metacommunity of bacteria, variance in microbiome composition across individual hosts is high (***Johnke et al., 2020***; ***Vega and Gore, 2017***; ***Taylor et al., 2022b***; ***Taylor and Vega, 2021***). While a characteristic worm microbiome can be defined at the family to genus level, notably featuring alpha- and gamma-Proteobacteria, Actinobacteria, Firmicutes, and Bacteroidetes (***Dirksen et al., 2020***), individual worm microbiomes are highly variable at the genus level and below (***Johnke et al., 2020***). Further, individual variation in bacterial load is substantial, with individual worms within a single experiment commonly containing 10^3^ to 10^5^ bacteria when colonized with communities of commensal microbes (***Dirksen et al., 2020***; ***Taylor and Vega, 2021***). This variation is highly repeatable run to run for worms of a given genotype but varies across genotypes (***Taylor and Vega, 2021, 2024***). This system therefore represents an ideal playground for examining the effects of host genotype and individuality on microbial interaction networks, and for determining the effects of heterogeneity on inference of these interactions from snapshot data.

Here we use snapshot microbiome data from large numbers of individual worms to infer drivers of microbiome community structure in *C. elegans*. We first confirm that the minimal microbiome used in these experiments has a recognizable ecology, with structural features similar to those found in microbiota of larger hosts. We then show that correlations in bacterial abundance across individual hosts are inconsistent with a near-neutral model without explicit interactions and that these correlations differ across host genotypes. Considering bacteria-bacteria and host-bacteria interactions as sources of (co)variance, we find that some statistical features of the data can be accounted for by allowing either inter-species bacterial interactions or correlated variation in model parameters across individual hosts. However, neither model was sufficient to explain all salient features of the data. Our results suggest that both microbe-microbe and host-microbe interactions shape the microbiome structure. At the same time, they illustrate fundamental difficulties in inference of interactions from snapshot data and suggest a need for improved models of host-microbiome dynamics.

## Results

### Ecology of a Minimal Worm Microbiome

The minimal eight-species worm microbiome community was originally described in (***Taylor and Vega, 2021***). Briefly, *C. elegans* adults were colonized on uniform mixtures of bacteria representing the native worm microbiome (***Dirksen et al., 2016***), and community composition data were obtained from dilution plating of worm intestinal contents onto solid agar where community members could be distinguished based on colony morphology. These data are, therefore, neither fixed-count nor compositional (we discuss the importance of this in *Appendix: Correlation Matrices*). Instead, absolute abundance of live, colony-forming bacteria was determined for each host in the data set, with an estimated threshold of detection around 1% relative abundance within each community. These data were produced by destructive sampling of individual worms and therefore represent a snapshot of community states across the population of hosts. As only one observation was possible for each individual, the data have no temporal auto-correlations. However, all worms within an experiment were synchronized on the same schedule before being exposed to the same meta-community of potential migrants under the same conditions, and thus plausibly represent samples from some shared underlying distribution(s) of states.

For these analyses, we focused on the best-represented host genotypes in the data set, each representing a minimum of 100 individuals: wild-type Bristol N2 (*n* = 153 worms over 10 independent experiments), MAPK immune-deficient AU37 (*n* = 104 worms, four experiments), and defecation mutant *exp-1(sa6)* II (*n* = 138 worms, five experiments) (Fig. 1). AU37 has loss of function in a *sek-1* MAP kinase kinase implicated in pathogen response (***Kim et al., 2002***). The defecation mutant *exp-1(sa6)* has an Exp (expulsion-defective) phenotype; this mutant has a more variable posterior-anterior contraction cycle than wild type (mean cycle length roughly 45 s for both; SD 3-6 s for wild type and 8-13 s in the mutant) and expels normally in roughly one out of every six defecation cycles (***Thomas, 1990***). We have chosen these particular mutants because both the immune response and the defecation variability are expected to affect the microbiome composition, providing additional signal for the analyses.

**Figure 1.**
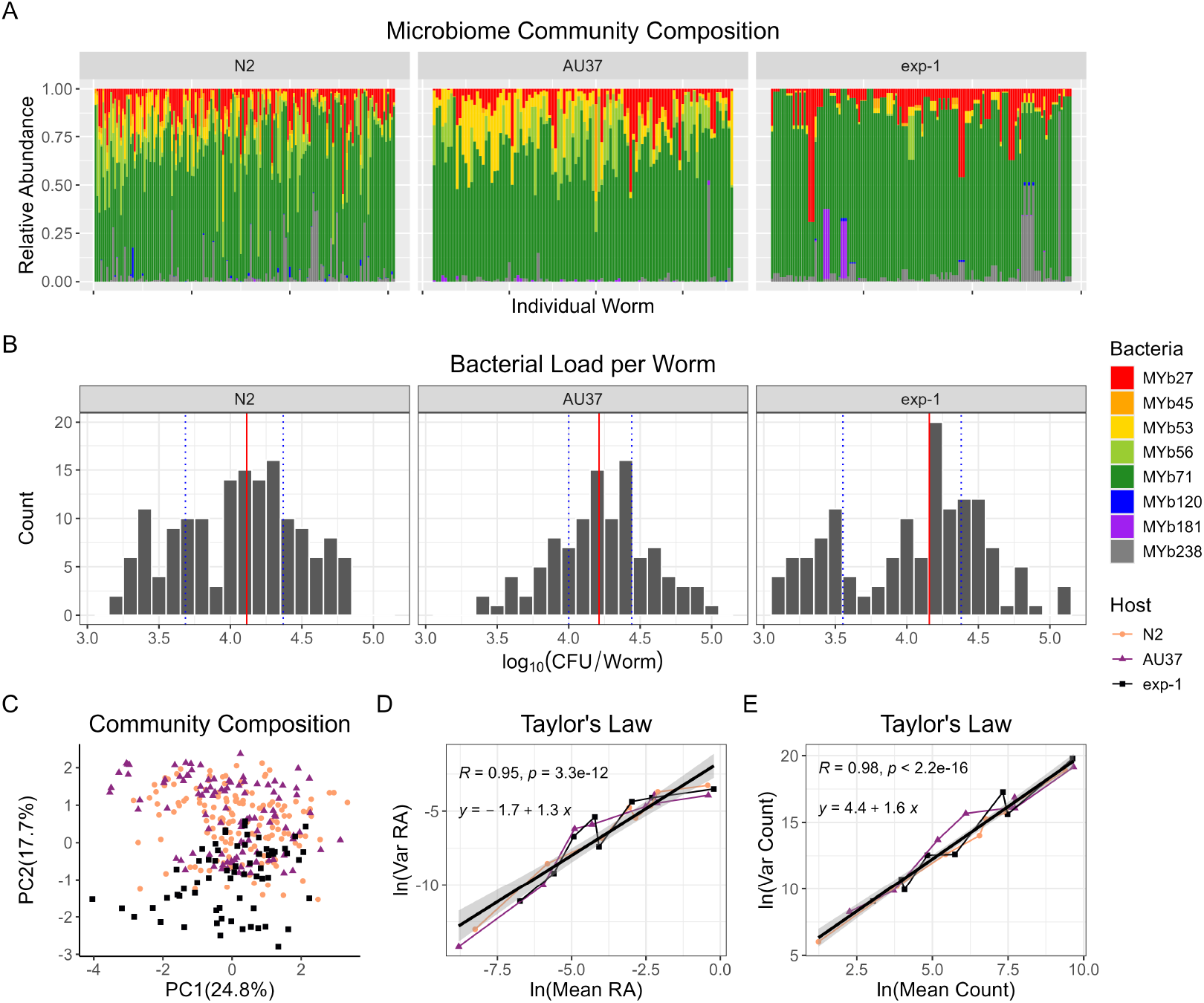
Ecology of *C. elegans* minimal microbiota. (A) Relative abundance of the eight bacterial taxa comprising worm intestinal communities within each host type. Each stacked bar represents one individual worm, with individuals sorted according to total bacterial load in the intestine (lowest to highest). (B) Distributions of total bacterial load in intestinal microbiome communities. Vertical solid line represents median density for that host type; dotted lines represent quartiles 0.25 and 0.75. (C) PCA of community composition (count) data. Each point represents one individual worm. (D-E) Taylor’s Law fits to (D) relative and (E) absolute abundance data for all eight community members; each point represents one bacterial taxon. Host genotype in (C-E) is indicated by symbol and color.

As previously described, adult worms colonized with this minimal eight-species microbiome of native gut bacteria (***Dirksen et al., 2016***) showed considerable individual heterogeneity in relative abundance of member taxa and in total bacterial load (***Taylor and Vega, 2021***). Despite variation within and between host types, we observed characteristic structure, such as a trend for high relative abundance of Proteobacteria (particularly *Ochrobactrum*) and lower relative abundance of Bacteroidetes (Fig. 1A), consistent with other observations in the worm (***Dirksen et al., 2020***; ***Johnke et al., 2020***). All three host types supported median bacterial loads just above 10^4^ colony forming units (CFU) per worm, with a range of approximately 10^3^ − 10^5^ CFU, although different host types had different distributions of bacterial load across this range. In particular, microbiome communities in the immune-compromised host AU37 had less variance and skewness and a smaller inter-quartile range than those in immune-competent hosts (Fig. 1 B). However, intestinal communities in wild-type N2 and MAPK immune-compromised AU37 were structurally similar, whereas the defecation mutant *exp-1* supported distinctive communities characterized mostly by lower relative abundance of *Bacillus* (MYb56) and *Rhodococcus* (MYb53) (Fig. 1 C). Further, the abundance distribution in the defecation mutant had heavier tails, with some individuals with very low or very high total bacterial load. Microbiome structures across all host types were broadly consistent with Taylor’s law (variance scaling as a power of the mean), with fits indicating *b* = 1.3 − 1.6 as the scaling exponent, depending on whether total or relative abundance data were used (Fig. 1 D-E), consistent with expectations for ecological community data.

### Space Limited Growth without Interactions

From these data, minimal microbiota in *C. elegans* could be characterized as ecological communities with distinctive structure, including characteristic patterns of variation across individual hosts. We therefore sought to determine the nature of the interactions responsible for the observed composition and structure of these communities. *A priori*, microbial communities within a host can be shaped by interactions with the host, including but not limited to environmental filtering of potential colonists, and by interactions among microbes in the space- and resource-limited environment provided by the host.

Before attempting to infer specific interactions, it is necessary to assess performance of a neutral model to determine whether the observed microbial population variability might have arisen purely due to noise. Therefore, we first developed a simple well-mixed stochastic model of microbial population dynamics, which considers only migration, birth, death, and neutral competition for space within the host:

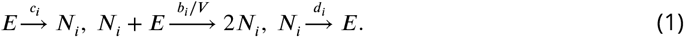

Here *E* represents an empty space and *N*_*i*_ a bacteria of type *i*. In this simple neutral model, all bacteria compete on equivalent terms; one bacterium of any type will occupy one empty space, and colonizing bacteria cannot be displaced through bacteria-bacteria interactions. In this set of “reactions”, a worm host is colonized by a bacteria of type *i* with a characteristic migration rate *c*_*i*_, after which bacteria undergo space-limited growth with a maximum doubling rate *b*_*i*_ and per-capita death with a constant rate *d*_*i*_. The sum of all the empty spaces and bacteria abundances within the gut is a constant, equal to the carrying capacity *V*. Note that rate constants are allowed to differ across bacterial taxa, but, for now, parameters are assumed to be the same in all worms of a given type. This model describes a simple scenario where bacterial taxon-specific colonization and growth determine the means and variances of the predicted population densities.

The mean field equations (see *Appendix: Space limited growth without interactions*) governing the means of the population densities *ϕ*_*i*_ are a generalization of the one used by (***Vega and Gore, 2017***):

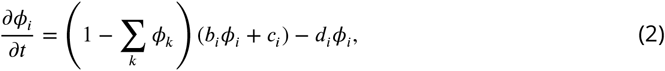

where *ϕ*_*i*_ is the abundance of the bacterial species *i* measured in the units of the carrying capacity, so that ∑_*i*_ *ϕ*_*i*_ ≤ 1.

The data in our experiment are snapshots of community composition rather than time series, as is often the case for microbiome data, and are thus not ideal for fitting population dynamics models (***Carr et al., 2019***). Further, fits of this model to the data are not unique, and many different parameter combinations will result in the same snapshot data. However, predictions about the expected (co)variances are more stable and can be used to assess suitability of the model without relying on precise parametric fits.

Using the system size expansion, in the *Appendix:Model Expansion*, we calculate the predicted covariances (not the correlation) for this simple model:

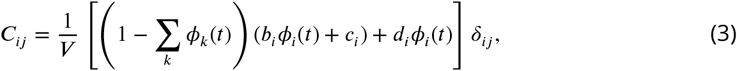

where *δ*_*ij*_ is the Kronecker delta function, *V* is the carrying capacity (volume) of the worm, and *ϕ*_*i*_(*t*) is the time-dependent ensemble-mean species abundance. This corresponds to a diagonal covariance matrix, such that there are no off-diagonal covariances (between bacterial taxa).

We tested this prediction by calculating the copula transformed correlation matrix (rankrank correlations) for each set of microbiome community data (***Nelsen, 2007***). We observed significant off-diagonal dependencies among bacterial taxa, inconsistent with expectations from the simple neutral model, Eq. (3), cf. Fig. 2. An alternate hypothesis to the no-interactions model is that niche-based interactions between bacteria will predominate. In general, most such inter-species interactions among bacteria are expected to be competitive (negative correlations) and weak (***Coyte et al., 2015***; ***Ho et al., 2024***), which would result in a sparse matrix of dependencies where most significant interactions are < 0. This is also inconsistent with the data, for all three host backgrounds.

**Figure 2.**
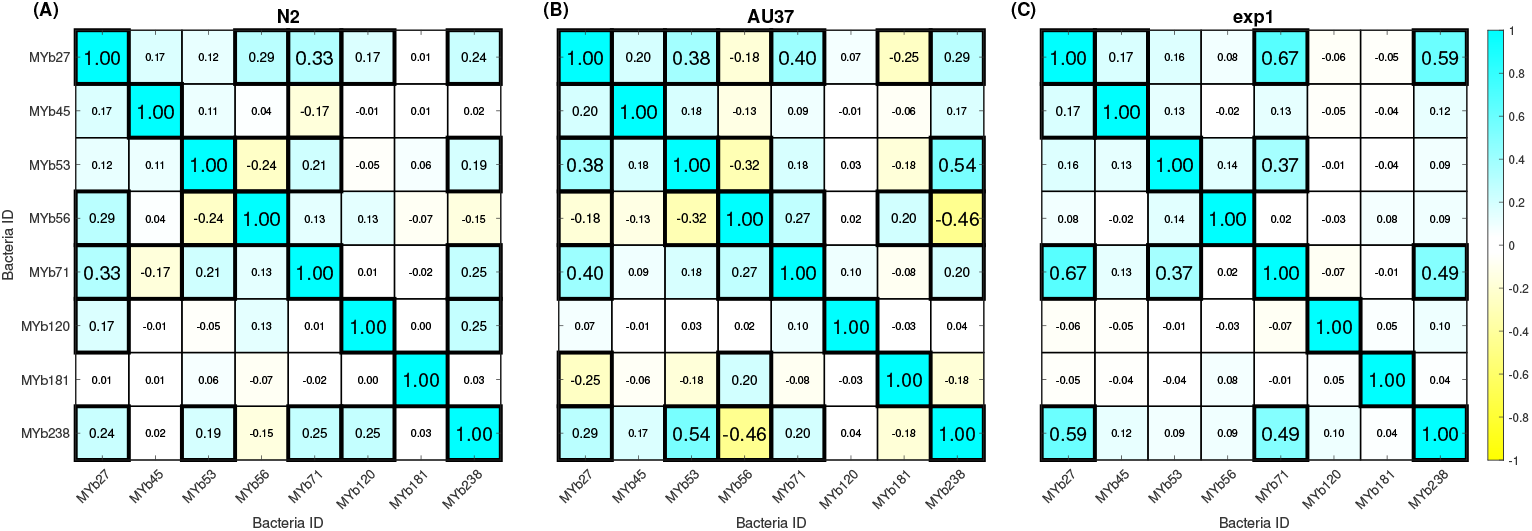
Correlation matrix for copula transformed community composition data (counts; absolute abundance) from (A) N2 (*n* = 153 worms), (B) AU37 (*n* = 104), and (C) *exp-1* (*n* = 138) hosts. Statistical significance for each correlation matrix was calculated by bootstrapping the copula transformed experimental data 1000X and re-calculating correlation matrices for each re-sample. Bonferroni correction was used to accommodate multiple hypothesis testing (with *n* = 28 possible off-diagonal correlations, family-wise confidence of 95% is obtained using a confidence of 99.8% for each correlation or a z-score of 3.12). Correlations with greater than 95% family-wise confidence are given a solid black border. The size of the text indicates the confidence of the entry being different than 0. Entries are colored by correlation value. Note that matrices are symmetric across the diagonal.

Indeed, in N2 hosts, there were 10 (out of 28 possible) statistically significant off-diagonal correlations. Most of these (8 of 10) were positive, and interactions were not evenly distributed across member taxa. For example, MYb27 (*Arthrobacter aurescens*) and MYb238 (*Stenotrophomonas*) were associated with all but one of these positive off-diagonal dependencies in the data set. Similar results were obtained for *exp-1* hosts, which showed no significant negative dependen-cies among bacterial taxa in copula-transformed data, while showing 5 statistically significant positive correlations, concentrated again in the same bacterial species. Although the N2 and AU37 community composition data were structurally similar (Fig. 1), AU37 host supported qualitatively different bacterial correlations: of 12 significant correlations, 4 were negative, and also of a larger magnitude than in the immune-competent host. Predictably, the same two bacterial species were responsible for 9 of the 12 significant interactions. Converting count data to relative abundance (see *Appendix:Correlation Matrices*) predictably resulted in a predominance of negative inferred dependencies, a well-known artifact due to the constraint that relative abundances must sum to 1.

Some inter-species correlations were roughly conserved across host types. Specifically, three bacterial taxa (MYb27, MYb71, and MYb238) showed significant positive dependencies in all three hosts. These dependencies were observed in absolute but not relative abundance. Consistent with this observation, total bacterial load in individual worms was positively correlated with counts of each of these bacterial taxa (see *Appendix: Community Composition Data*). Origins of negative dependencies were more variable. Bacteria MYb53 (*Rhodococcus*) and MYb56 (*Bacillus*) showed a negative dependency conserved across N2 and AU37 hosts which was not obviously dependent on total bacterial load (see *Appendix: Community Composition Data*). Negative correlations appearing in AU37 but not in either immune-competent host (MYb56/MYb238, MYb56/MYb27) were associated with relatively high rates of reciprocal absence of these taxa in AU37 hosts (see *Appendix: Community Composition Data*).

These results indicated that the simple neutral model with constant parameters was insufficient to match the correlation structure in these data. Further, the observed correlation matrices provided information on possible differences in control of microbiome communities across host types. In immune-competent (N2 and *exp-1*) hosts, counts of different bacterial taxa tended to be positively correlated with one another and with total bacterial load in individual worms (see *Appendix*). This suggests that these communities may not be structured primarily by inter-species interactions among bacteria. Instead, these results suggest that individual immune-competent hosts may be more or less permissive for sets of bacterial taxa, such that counts of multiple taxa rise and fall according to the host state. This is consistent with earlier conclusions from a different analysis of the same data set (***Taylor and Vega, 2021***). However, negative dependencies tended to be specific to pairs of bacterial taxa, which could (but does not necessarily) indicate negative inter-species interactions. Further, facilitation among bacteria (e.g., cross-feeding) can produce positive correlations between taxa and increase overall community yield. While the number of facilitative interactions needed to explain these data would be large, the possibility cannot be ruled out entirely.

### Bacterial interactions

While off-diagonal correlations between bacterial taxa might arise from direct bacteria-bacteria interactions, differences in the environment presented by individual hosts, or both, the most obvious way to obtain off-diagonal correlations is to allow inter-species interactions among bacteria. This is canonically expressed in the gLV model as a matrix of pairwise interactions *A*, where each term *A*_*ij*_ indicates the per-capita effect of species *j* on growth of species *i* (***Jones and Carlson, 2019***; ***Joseph et al., 2020***). However, as indicated earlier, snapshot data are not ideal for inference of these population-dynamic parameters. Thus, in what follows, we chose to *not* quantitatively fit the abundance data to a gLV model, but to focus on qualitative predictions of such models.

In constructing the model, we needed to account for the stochasticity of the demographic processes. Thus, we chose to analyze an extension of the stochastic model, Eq. (1), which includes bacterial interactions. While similar to the gLV models, the mean-field equations of this stochastic model do not map onto the gLV structure exactly. Specifically, we constructed a model with no spatial structure. We broadly classified bacteria-bacteria interactions into three categories by sign: warfare (decrease both species number by one), predation / competition (increase one species while decreasing the other species), and mutualism/facilitation (increase both species by one). These reactions are of the lowest interaction order (pairwise) that is biologically relevant, and we neglected higher order interactions for simplicity. Overall, the corresponding “reaction” terms in this model are:

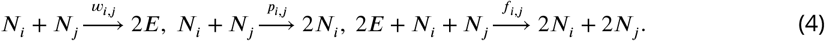

As shown in *Appendix: Bacterial Interactions*, the resulting covariances (not correlations!) for *i* ≠ *j* within the model are:

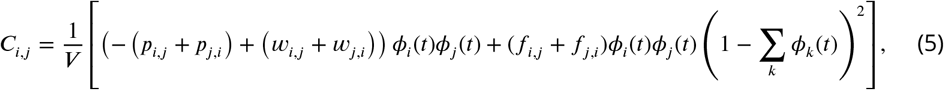

where *ϕ*_*i*_(*t*), again, are the ensemble-mean abundances of the species in the units of the carrying capacity, which solve the mean field equations for the model in the *Appendix*. For *i* = *j*, the diagonal covariances are:

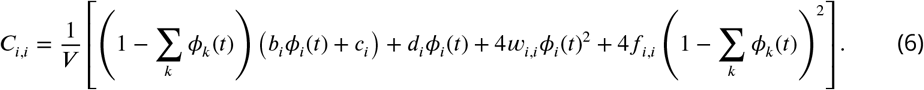

By inspecting these equations, we notice that the three different types of interactions, parameterized by *w, p, f*, produce different patterns in the covariance matrices. Predation / competition produces negative off-diagonal entries but does not contribute to the on-diagonal variance, while both warfare and facilitation produce positive off-diagonal entries and contribute positively to the on-diagonal variance. Unlike warfare, which releases empty space and can therefore continue in saturated communities, facilitation is modulated by the need for empty sites for growth.

We cannot fit a model with specific bacteria-bacteria interactions to our data: this would be over-parameterized, as there are only 8 independent species means and 32 off- and on-diagonal species covariances, while there are 28×3=84 parameters describing the three types of interactions for all pairs of species. There are, however, some qualitative predictions from Eqs. (22, 23) that can be made without fitting. Specifically, the presence of significant negative off-diagonal correlations in bacterial abundance data (such as in Fig. 2) can only be explained by predation / competition interactions within this model. Positive correlations could be produced by facilitation and/or warfare, which are difficult to disambiguate from snapshot data. The off-diagonal covariance elements are predicted to scale as the product of the mean abundances. If there is no facilitation, then the ratio of the off-diagonal covariances to their means 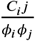 should remain constant with time. We test this in the *Appendix: Bacterial Interactions* and find that the ratio changes slightly (but statistically significantly) with time indicating that facilitation would be necessary.

However, while bacteria-bacteria interactions are a plausible sources of off-diagonal terms, it is not clear that such interactions are either sufficient or necessary to explain the data. First, the inferred dependencies among bacterial taxa were not conserved across different host types, cf. Fig. 2, suggesting that, if we insist to use Eq. (20) to model the data, the interactions between the species must be host-specific, which is not easily explainable. This would be particularly problematic for N2 and AU37 hosts, which supported structurally similar communities covering a similar range of population densities. Thus the model, Eq. (20), may not be sufficient for explaining the data.

At the same time, the model may also be not necessary to explain the correlations since, as detailed in the next section, an alternate model without bacteria-bacteria interactions, but with differences between hosts, may have similar explanatory powers.

### Host Heterogeneity

Differences between individual hosts can also produce correlations in microbiome data ***Armitage and Jones (2019***). “Hidden” heterogeneity within synchronized, isogenic populations of *C. elegans* is associated with individual differences in development (***Natesan et al., 2023***), stress response (***Vertti-Quintero et al., 2021***; ***Rea et al., 2005***), and lifespan (***Suda et al., 2009***; ***Wu et al., 2006***). It is plausible that this heterogeneity could also affect interactions with commensal bacteria. In this case, individual hosts might pull from different parameter ranges for bacterial growth in the intestine, for example due to differences in innate immune function or nutrient provisioning. If parameters within a host are correlated across bacterial taxa, positive correlations can arise simply due to fast-growing species growing quickly together. It is likewise plausible that negative correlations could arise due to differences between hosts due to parameter anti-correlations, for example if an immune response that suppresses one bacterial taxon precludes activation of responses that are effective against a second (***Yan et al., 2020***; ***Zhang et al., 2021***). Ecologically, this would produce a sort of Moran effect (***Cazelles and Boudjema, 2001***), where correlations among non-interacting units emerge due to correlations between environmental variables. Such host-induced correlations, however, will not be a signature of bacteria-bacteria interactions.

We, therefore, sought to determine whether the observed off-diagonal correlations could arise due to differences between hosts, and if so, under what conditions and constraints in parameter space. It is possible to calculate analytically how the covariance matrix calculated from bacterial count data depends on the covariance matrix of the parameters of the meanfield model if we assume that these parameters are drawn independently from Gaussians centered at their best fit values. This assumption is almost certainly untrue (see *Appendix: Host Heterogeneity*), but as we are attempting to recapitulate the range of plausible signs rather than the actual covariances from the data, it is sufficient for our purpose. To the lowest order, the covariance of the data *C*_data_ is related to the covariance of the parameters *C*_param_ by *C*_data_ = *G*^*T*^ *C*_param_*G* where 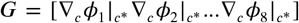 and *c*^*^ is the set of parameters resulting in the best fit. We can invert this relationship by using pseudo-inverses calculated by using SVD since, in general, *G* is a singular matrix (*G*^+^ = *G*^*T*^ (*GG*^*T*^)^−1^). This gives a non-unique solution with

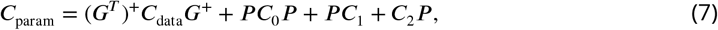

where *P* = *I* −*GG*^+^ = *I* −(*G*^*T*^)^+^*G*^*T*^ = *P* ^*T*^ is the null space projection operator of *G*, and *C*_0_, *C*_1_, *C*_2_ are constant matrices. This relationship for *C*_param_ is only unique when the null space projector is zero, *P* = 0. However, with a L2 norm regularizer encouraging correlation coefficients to be as small as possible, we achieve a unique solution

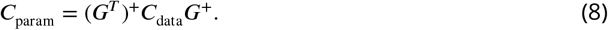

Thus, to obtain parameter correlations from data, we first numerically calculated the gradient of the population densities of bacteria w. r. t. parameter values, evaluated using the best fit parameters. We then numerically followed this pseudo-inverse procedure to find the covariance and correlation structure of the parameters.

To calculate the dependence of the population densities on the parameters, here we worked with a version of the simple neutral model, where bacterial birth rates *b* and colonization rates *c* are drawn from the same distribution for all bacterial taxa, species-specific death rates *d*_*i*_ are drawn from different distributions, and capacity *V* is held constant as before. This model does not include pairwise interactions among bacteria. Thus any apparent correlations between taxa result, in this model, from parameter correlations rather than direct bacteria-bacteria interactions. This form of the model was selected to minimize the total number of parameters while remaining biologically interpretable (see *Abstract*).

We found that both positive and negative correlations among bacterial taxa could be produced by allowing parameters to co-vary across hosts in the absence of explicit bacteria-bacteria interactions. Positive correlations among parameters were common, particularly in N2 data, consistent with the predominance of positive correlations in this data set (Fig. 2). The AU37 data set produced more negative parameter correlations, as expected.

We verified these fits by running deterministic simulations with parameters drawn from Gaussians centered around the best fit parameters and their predicted covariances, to simulate the effects of this distribution of heterogeneity across hosts. Here each simulation run represents one worm drawn from the parameter space of the fitted model. Numerical experiments of the simple neutral model (both numerical integration of mean field equations and stochastic Gillespie simulations (***Gillespie, 2001, Gillespie 1977***)), where we forced the parameters to be independent and drawn from Gaussians centered around their best fit values, produced small positive and negative off-diagonal covariances but very close to zero, as expected (SI FIG. 10). By contrast, simulations with parameter covariances as in Fig. 3A-C generated both positive and negative abundance covariances, qualitatively similar to observations from our experimental data. However, in general the simulated correlations were smaller than those in the real data, Fig. 3D,E, though the difference was pretty small for stochastic simulations. This suggests that parameter distributions in real data may have longer, non-Gaussian tails.

**Figure 3.**
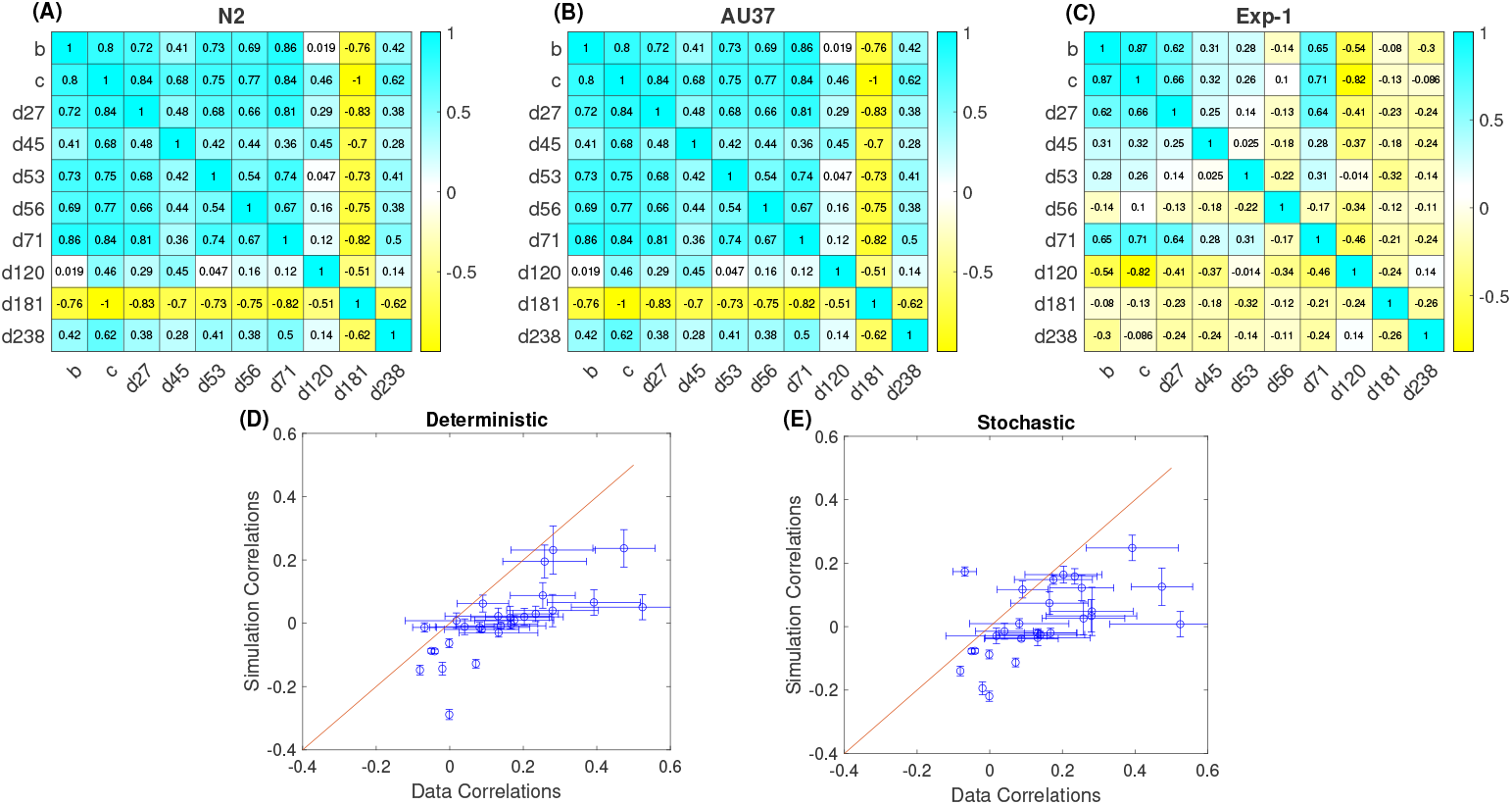
The inferred correlation matrix of the model parameters for (A) N2, (B) AU37, and (C) *exp-1* worms calculated by using the pseudo inverse method acting on the empirical correlation matrix of microbial abundances. (D) Comparison of empirical and simulated microbial abundance correlations for the N2 worm. For simulated correlations, parameter values were sampled from the matrix in (A), and deterministic population dynamics was assumed. (E) Same as (D), but for stochastic population dynamics.

## Discussion

Here we used data from a tractable host-microbiome model, the nematode *C. elegans*, to explore the structure of variation in the minimal microbiome, to understand which effects contribute to the microbiome composition, and to elucidate difficulties in inferring these effects from the microbial community abundance data. These minimal host-associated communities show characteristic structure with definable differences between and heterogeneity within host types. They conform to known ecological laws, suggesting a host-dependent ecology shaped by some consistent set(s) of interactions. This has been observed for other minimal host-associated microbiota (***Shetty et al., 2022***), suggesting the utility of these simplified model systems for studies of microbiome ecology.

The microbial count-based snapshot data in our experiments are structurally quite different from standard microbiome metagenomic data sets. Indeed, whereas metagenomic data from undefined, species-rich assemblages tend to be sparse (abundant zeros), long (many more taxa than samples), and compositional (***Joseph et al., 2020***), our data are dense (relatively few zeros; high prevalence of individual taxa), wide (the same eight taxa in order 10^2^ samples), count-based rather than proportional, and not fixed-count. Further, in our data, individual hosts are destructively sampled and only appear once in the data set. Such single observations are not atypical in other microbiome data, although there has been a recent trend toward collection of time series (***Caporaso et al., 2011***; ***Ji et al., 2019***; ***Inamine et al., 2018***; ***Gerber et al., 2012***; ***Poyet et al., 2019***). These differences mean that standard approaches to analysis of microbiome community data are not applicable to our data, and, conversely, that some techniques inappropriate for sparse compositional data can be applied. Yet, we show that fundamental problems of inference are conserved for these count-based microbiome data, and we exploit the non-canonical structure of these data to illustrate fundamental difficulties in determining interactions within microbiomes.

We found that a simple neutral model is inconsistent with data for microbiome communities in *C. elegans*. Colonization of the worm intestine is a stochastic process, and the rate, at which migrants are introduced, can drive distributions of community composition when bacterial strains interact more or less neutrally (***Vega and Gore, 2017***). The assumption of neutral interactions is rarely literally true, and biotic interactions among colonizing bacteria can shape worm intestinal communities (***Johnke et al., 2020***), suggesting that these interactions should not be ignored. However, when specific bacteria-bacteria interactions are not known *a priori*, our results indicate that signals from heterogeneous host states cannot easily be distinguished from those produced by interactions among bacteria, even when absolute abundance is known.

Host-dependent parameter variability is a specific example of so called latent variables models, which have become popular in other quantitative biology fields, such as neuroscience, immunology, or community ecology ***Köster et al. (2014***); ***Warton et al. (2015***); ***Desponds et al. (2016***); ***Morrell et al. (2024***). In principle, latent variability results in very specific signatures in data (***Schwab et al., 2014***; ***Aitchison et al., 2016***), which should allow one to distinguish it from specific interactions. However, this requires very large systems (in our case, microbial communities with a lot of species) and sample sizes that scale with the system size (***Ngampruetikorn et al., 2023***), which is not easy to generate in a system such as ours. In contrast, most more traditional methods for inference of interactions in the context of systems biology (***Natale et al., 2017***) get confused by latent variability effects, or also require unrealistically large system and sample sizes to achieve high statistical power. It thus seems that distinguishing effects of the host and of bacterial-bacterial interactions will likely require different experimental setups, rather than more sophisticated analysis tools.

Interestingly, while distinguishing host-bacteria and bacteria-bacteria interactions has proven to be hard, neither of these interactions were structurally sufficient to explain the salient features of our data. We observed that correlations between bacterial species abundances were not completely unstructured, with some taxa associated with positive vs. negative inter-species correlations, but that this was far from being a general rule. Further, we observed that bacterial load varied substantially between individual hosts, and that the proportional composition of these communities was largely unaffected by total size. Bacteria-bacteria interactions can be uncorrelated between pairs and asymmetric within a pair, such that the resulting covariances among species can vary freely, consistent with the weak structure observed in correlation matrices. However, these models do not allow total population size to vary significantly. By contrast, allowing parameter correlations (correlated host heterogeneity) can produce differences in effective population size, but the resulting correlation matrices are highly structured.

Indeed, the wide range of total bacterial loads is difficult to explain without allowing alternate states in these host-microbiome systems. The nature of those states remains unclear. First, it is unclear which parameter(s) are or should be associated with alternate host-microbiome states. Further, as each individual host is represented only once in a data set, it is not clear whether individual worms should have static positions in this hypothetical state space. Previous work in environmental systems (***Martin-Platero et al., 2018***; ***Shade et al., 2014***) and in human microbiomes (***Zaoli and Grilli, 2022***) has suggested evidence for state-switching, where the carrying capacity of sub-sets of community members appears to change abruptly in time series data. However, *C. elegans* is a short-lived, anatomically simple host, which is held under constant conditions in our experiments. It is plausible that a worm and its microbiome are static in ways that other systems are not. Further work is required to address the question of host states and the corresponding microbial load variability. More broadly, these results support, in principle, the recommendation for determination of bacterial load per sample as part of best practice in microbiome community data collection (***Lloréns-Rico et al., 2021***).

Although our data do not allow specification of unique population dynamics models, the differences we observed between microbiota in hosts of different types provide potential insight into sources of variation in host-associated microbiomes. For example, the *exp-1* mutant has futile defecation cycling, expelling gut contents on average once every six cycles, and the variance in cycle length is larger than for the N2 wild-type (***Thomas, 1990***). This suggests a scenario where irregular, large gut motility events act as disruptions to the microbiome, resulting in increased variance in microbial load and altered community composition (Fig. 1). Similar effects have been observed in the zebrafish microbiome (***Wiles et al., 2016***; ***Schlomann et al., 2019***), although the mechanics are likely to be different due to the anatomy of the gut in these systems. By contrast, communities in N2 and the MAPK-deficient AU37 host were compositionally similar, but AU37-associated communities were enriched in negative bacteria-bacteria correlations (Fig. 2). Under the assumption that apparent interactions are an artifact of parameter correlations, these data required extensive negative co-variances among parameters in only this host (Fig. 3), which is difficult to support biologically. Instead, this suggests that communities in AU37 are shaped less strongly by host control; a similar hypothesis has been put forth for commensal communities in worms deficient in activity of the central IGF-1 transcription factor *daf-16*, which show a similar but more marked trend in bacteria-bacteria correlations as compared with N2 wild type (***Taylor and Vega, 2021***; ***Zhang et al., 2021***).

While these results suggest that microbiome communities in *C. elegans* are the product of non-neutral underlying dynamics, it remains unclear which interactions are primarily responsible for shaping these communities. These and prior results suggest an important role for host control of the microbiome, but the dynamics imposed by the host are not well understood. It is apparent that high-quality, abundance-based time series data will be necessary to address this problem, and we suggest that simple model systems of this type are uniquely suited to investigate these questions.

## Methods

### Strains and Culture Conditions

Arthrobacter aurescens (MYb27), Microbacterium oxydans (MYb45), Rhodococcus erythropolis PR4 (MYb53), Bacillus sp. SG20 (MYb56), Ochrobactrum vermis (MYb71), Chryseobacterium sp. CHNTR56 (MYb120), Sphingobacterium faecium (MYb181), and Stenotrophomonas sp. (MYb238) were obtained from the Schulenberg lab (***Dirksen et al., 2016***). *C. elegans* S medium and M9 worm buffer were prepared according to standard WormBook protocols (***Stiernagle, 1999***). M9 worm buffer + 0.1% Triton X-100 (M9TX01) was used in some protocols to prevent worms from sticking to plastic during handling and pipetting (***Taylor et al., 2022b***). *C. elegans* axenic medium (AXN) was prepared according to published protocols (***Houthoofd et al., 2002***) by autoclaving 3g yeast extract and 3g soy peptone (Bacto) in 90 ml water, and subsequently adding 1g dextrose, 200*μ*l of 5 mg/ml cholesterol in ethanol, and 10 ml of 0.5% w/v hemoglobin in 1 mM NaOH before filling to 100 ml. Salt-free nutrient agar [FORMULA] was used for plating and enumeration of live bacteria.

Laboratory wild-type N2 Bristol, *exp-1(sa6)* II (CGC JT6) (***Thomas, 1990***), and AU37 *glp-4(bn2)* I; *sek-1(km4)* X *C. elegans* were obtained from the Caenorhabditis Genetic Center, which is funded by NIH Office of Research Infrastructure Programs (P40 OD010440). Nematodes were grown, maintained, and manipulated using standard techniques (***Stiernagle, 1999***; ***Taylor et al., 2022b***). Briefly, breeding stocks were maintained on NGM plates + OP50 at 25°C (16 °C for temperature-sensitive strain AU37 and synchronized using a standard bleach/NaOH protocol where eggs were allowed to hatch out in M9 worm buffer overnight (16h) with shaking (200 RPM) at 25°C. Starved L1 larvae were transferred to 10cm NGM plates containing lawns of *E. coli* expressing *pos-1* RNAi (***Kamath et al., 2001***) and incubated at 25°C for three days to produce reproductively sterile adults; temperature-sensitive sterile AU37 were grown to adulthood on OP50 under the same conditions. Worms were then transferred to liquid S medium + 200 *μ*g/ml gentamicin + 50 *μ*g/ml chloramphenicol + 2X heat-killed OP50 (to trigger feeding) for 24 hours, resulting in largely germ-free adults (***Taylor and Vega, 2021***). Adult worms were washed via sucrose floatation, then rinsed 3X in 10 mL M9TX01 and once in 10 mL S medium to remove sucrose, before colonization.

### Worm Colonization and Bacterial Community Counts

Colonization of adult worms with an eight-species minimal microbiome community was carried out as in (***Taylor and Vega, 2021***). Briefly, bacterial strains were grown individually from glycerol in 1 mL LB for 48 hours at 25°C with shaking at 200 RPM to reach stationary phase. Cultures were then diluted to 10^8^ CFU/ml in 1ml S medium + 1% AXN and centrifuged 2 minutes at 10K RPM [IN G] to pellet. Supernatant was removed, and pellets were resuspended in 1ml fresh S-medium + 1% AXN. Communities were created by mixing equal volumes of all eight bacteria,

Colonization was performed in well-mixed liquid media according to standard protocols for this lab (***Taylor and Vega, 2021***; ***Taylor et al., 2022b***) to ensure that all individual worms experienced a uniform environment and had equal access to all potential colonists, for the duration of colonization. Germ-free adult worms were resuspended in S-medium + 1% AXN to a concentration of 1000 worms/ml. Aliquots of 180*μ*l were pipetted into 96-well deep culture plates (1.2ml well volume, VWR). 20 μl of each bacterial suspension was added to each well (final volume 200 *μ*L). Plates were covered with Breathe Easy sealing membranes and incubated with shaking at 200 RPM at 25°C.

After two days, worms were washed to remove external bacteria and re-fed on a freshly assembled metacommunity. This was done to enforce an evenly distributed community, maximize viability of potential colonists, and minimize bacterial evolution that might lead to unpredictable divergence across replicates. In this step, worms were washed twice in 1ml of M9 Worm Buffer + 0.1% v/v Triton X-100 (M9TX1), rinsed once with S-medium + 1% AXN to remove surfactant, and resuspended in 180 μl of S-medium + 1% AXN in wells of a fresh 1.2 mL deep 96 well plate before addition of 20 μl fresh bacteria.

After four days total colonization, worms were washed 2x in 1 mL M9TX01 to remove the bulk of external bacteria, then surface bleached and mechanically disrupted according to standard protocols for the lab (***Taylor et al., 2022b***). The resulting bacterial suspensions were dilution plated in 100 *μ* L aliquots onto 10cm nutrient agar (NA) plates and incubated for 48-72 hours at 25°C before colony counting to determine numbers of each bacterial species within individual worms.

## Data Availability

Data and code are available from Emory Dataverse (https://doi.org/10.15139/S3/NV1OIV).

## Appendix 1

### Correlation matrices

For completeness, here we calculate the Pearson correlation matrices for N2, AU37, and *exp-1* hosts, using both the raw count data (Appendix Fig. 1) and the copula transformed relative abundance data (Appendix Fig. 2) (note that Fig. 2 in the *Main Text* is correlations of the copula transformed community composition data). In general, it is **inadvisable** to calculate correlations on relative abundances. This is because relative abundances are defined such that the sum of all the abundances sum to one, ∑_*i*_ *r*_*i*_ = 1. This added constraint induces correlations that are not present in the raw data. If the absolute abundance of one species goes up the relative abundances of all other species will go down to compensate. The relative abundance *r*_*i*_ can be completely determined from the relative abundances of the other species *r*_*i*_ = 1 − ∑_*k*≠*i*_ *r*_*k*_. Mathematically we find that the covariances of relative abundances are related such that a particular covariance is equal to the negative sum of all other covariances in that column or row:

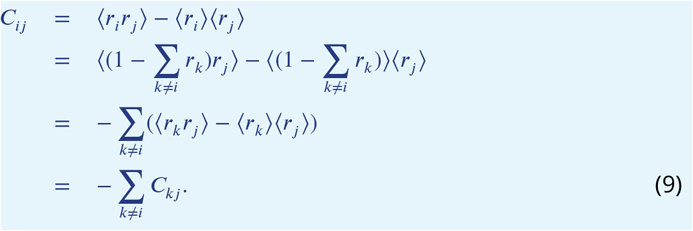

Using this result, it is clear that the correlations of relative abundances are similarly related:

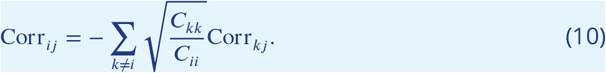

This relationship in correlation of the relative abundances is dramatically displayed in Appendix Fig. 2 in MYb71.

**Appendix 1—figure 1.**
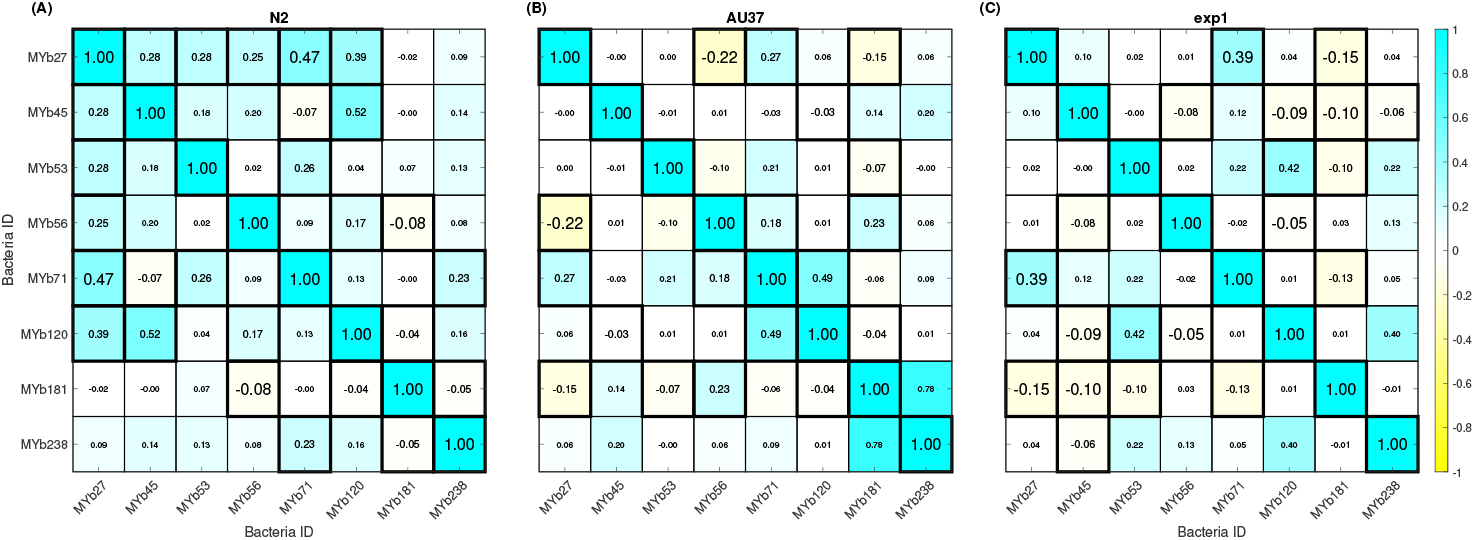
(Pearson) correlation matrix for raw community composition data (counts; absolute abundance) from (A) N2 (*n* = 153 worms), (B) AU37 (*n* = 104), and (C) *exp-1* (*n* = 138) hosts. Signal to noise ratio for each correlation matrix was calculated by bootstrapping the raw data 1000X and re-calculating correlation matrices for each re-sample. Bonferroni correction was used to accommodate multiple hypothesis testing (with *n* = 28 correlations, family-wise confidence of 95% is obtained using a confidence of 99.8% for each correlation or a z-score of 3.12). Correlations with greater than 95% family-wise confidence are given a solid black border. The size of the text indicates the confidence of the entry being different than 0. Entries are colored by correlation value. Note that matrices are symmetric across the diagonal.

**Appendix 1—figure 2.**
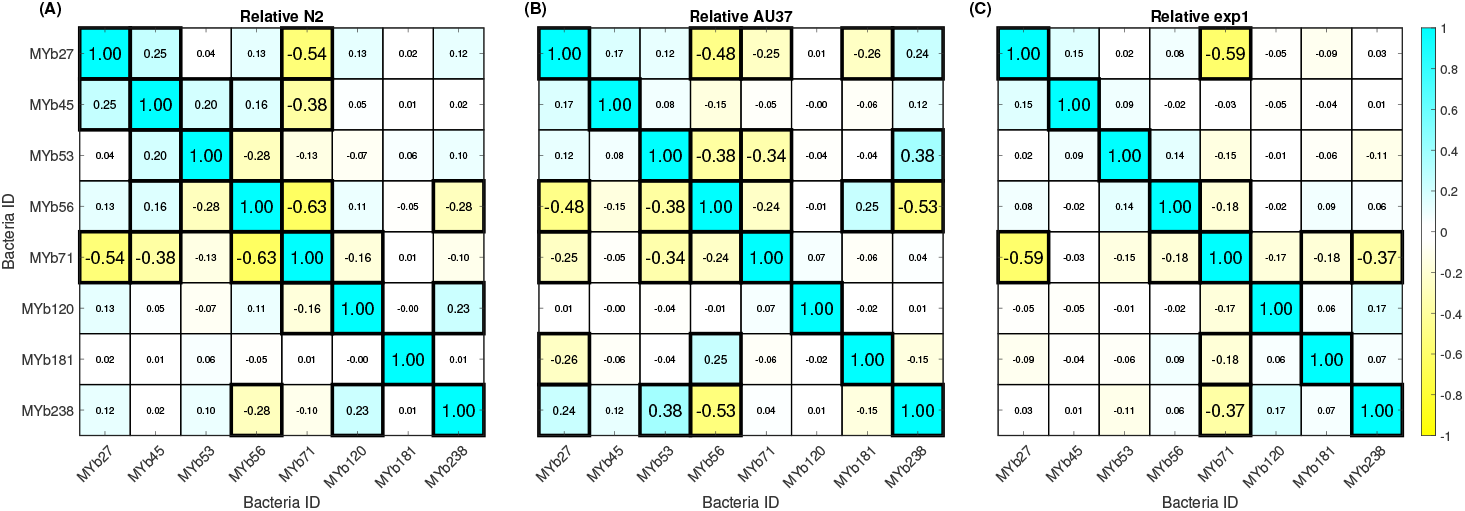
Correlation matrix for copula transformed relative community composition data (counts) from (A) N2 (*n* = 153 worms), (B) AU37 (*n* = 104), and (C) *exp-1* (*n* = 138) hosts. Plotting conventions same as in Appendix Fig. 1.

#### Community Composition Data

To further explore the statistical structure of our dataset, we calculated relationships between bacterial load/population size and various metrics of community composition and correlation. Overall, abundance of individual bacterial taxa tended to increase with total bacterial load (Appendix Fig. 3). This relationship was strongest for the highly prevalent and abundant bacterial taxon MYb71, which was frequently the most abundant taxon within individual worms. Other highly abundant and prevalent taxa (MYb27, MYb238) likewise showed a strong correlation to total load, with slopes at or near 1. MYb181, the least prevalent taxon in the data set, showed the weakest correlation with total bacterial load.

**Appendix 1—figure 3.**
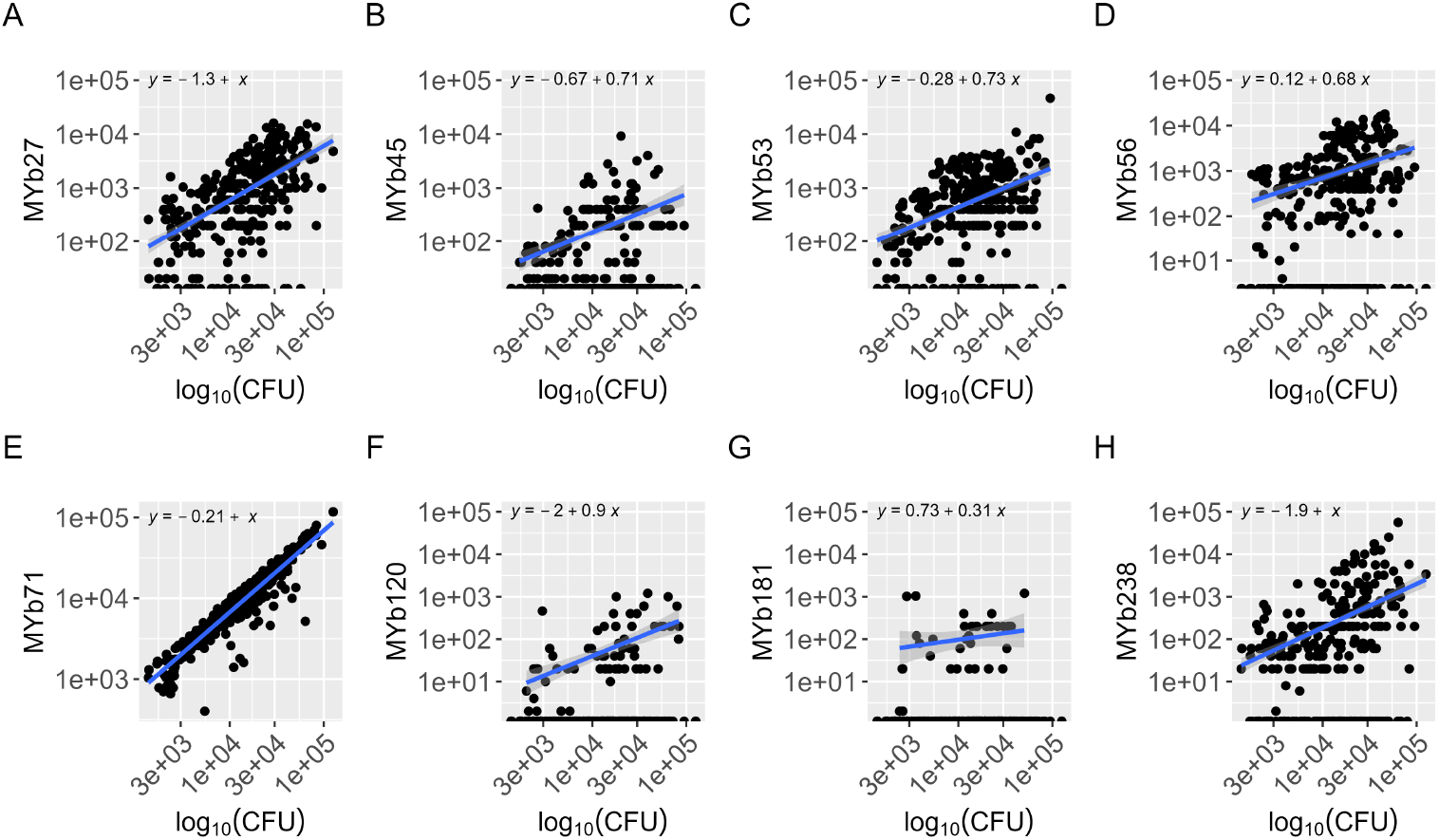
Abundance of individual bacterial taxa vs. total bacterial load in single worms. Data are pooled across all three host types.

Relationship between population size of individual bacterial taxa and total bacterial load were consistent with inferred dependencies between bacteria (see *Main Text*) (Appendix Fig. 4). Bacterial taxa that were individually strongly correlated with total bacterial load were inferred to have significant positive dependencies across all three host types, suggesting that the dependencies between these bacterial taxa could be explained by correlations with bacterial load in individual hosts rather than direct interactions between bacterial taxa. The negative dependency between MYb53 (*Rhodococcus*) and MYb56 (*Bacillus*) conserved across N2 and AU37 hosts was not as obviously explained by underlying dependency on total bacterial load in the intestine, suggesting that a different underlying mechanisms is required. A direct interaction between bacterial taxa is plausible; for example, while these taxa are not closely related, both are Gram-positive aerobes and may therefore be in competition for some specific niche. Note that MYb53 and MYb56 were not significantly dependent in data from the *exp-1* host, where information on this pair is limited due to the very low abundance and prevalence of MYb56 in these data. Negative correlations appearing in AU37 but not in either immune-competent host (MYb56/MYb238, MYb56/MYb27) were associated with relatively high rates of reciprocal absence of these taxa in AU37 hosts (Appendix Fig. 5).

**Appendix 1—figure 4.**
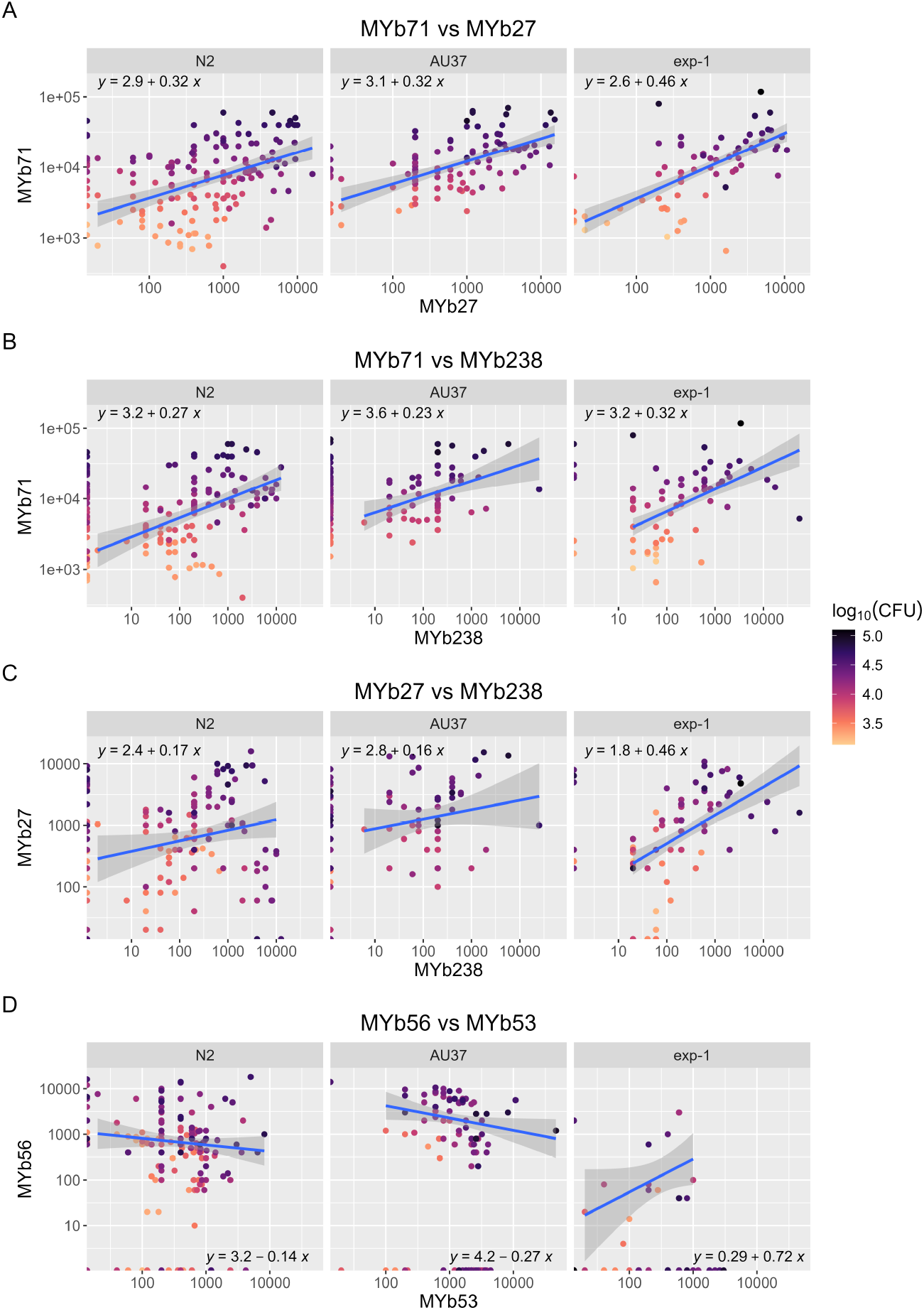
Dependencies among bacterial taxa that are conserved across host types. (A-C) Positive dependencies conserved across all three host types generally reflect underlying correlations with total bacterial load per worm; (D) A negative dependency conserved across N2 and AU37 hosts reflects a negative correlation between counts of two bacterial taxa. Each data point represents one individual worm.

**Appendix 1—figure 5.**
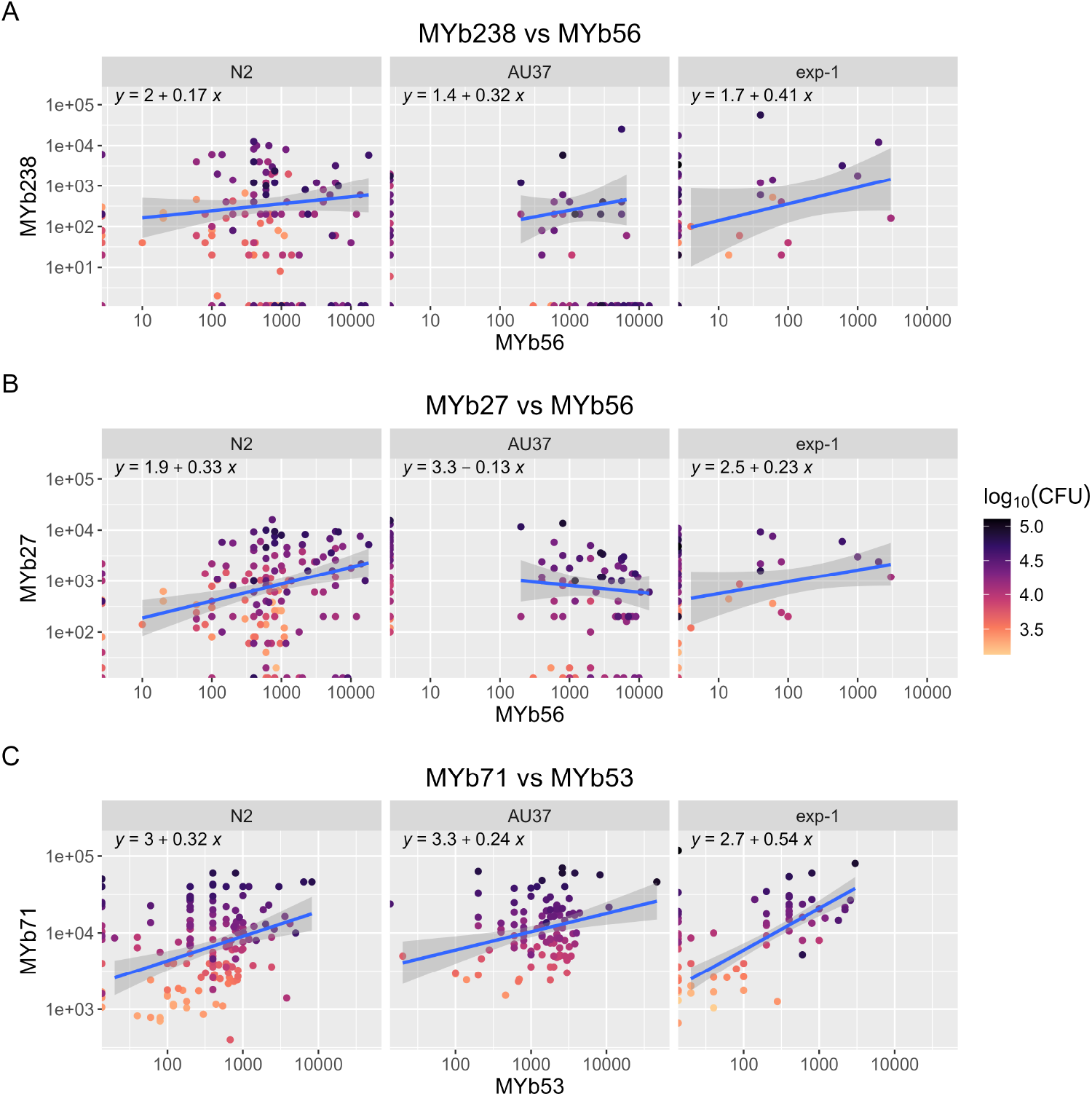
Dependencies not conserved between AU37 and immune-competent hosts. (A-B) Pairs showing negative dependency in copula-transformed AU37 data but not in data from N2 or *exp-1* hosts have high rates of reciprocal absence in AU37. (C) A significant positive dependency in immune-competent hosts is present as a trend in AU37 hosts. Each data point represents one individual worm.

While community richness and bacterial load were positively correlated in AU37 and *exp-1* but not N2 hosts, other measures of diversity (Simpson, Shannon (***Haegeman et al., 2013***)) were negatively correlated with bacterial load in N2 but not in other hosts (Appendix Fig. 6). This suggested that larger bacterial communities were associated with higher prevalence of rare taxa in AU37 and *exp-1*. Plotting prevalence of each bacterial species vs. total bacterial load indicated that this was essentially correct, as prevalence of several taxa in AU37 and *exp-1* data sets (MYb45, MYb56, MYb120) increased with total bacterial load. Of these, only MYb120 increased in prevalence in N2 hosts as bacterial load increased. One of these taxa (MYb45) instead decreased with bacterial load in N2 hosts, and MYb56 had generally lower prevalence and relative abundance in non-N2 hosts.

**Appendix 1—figure 6.**
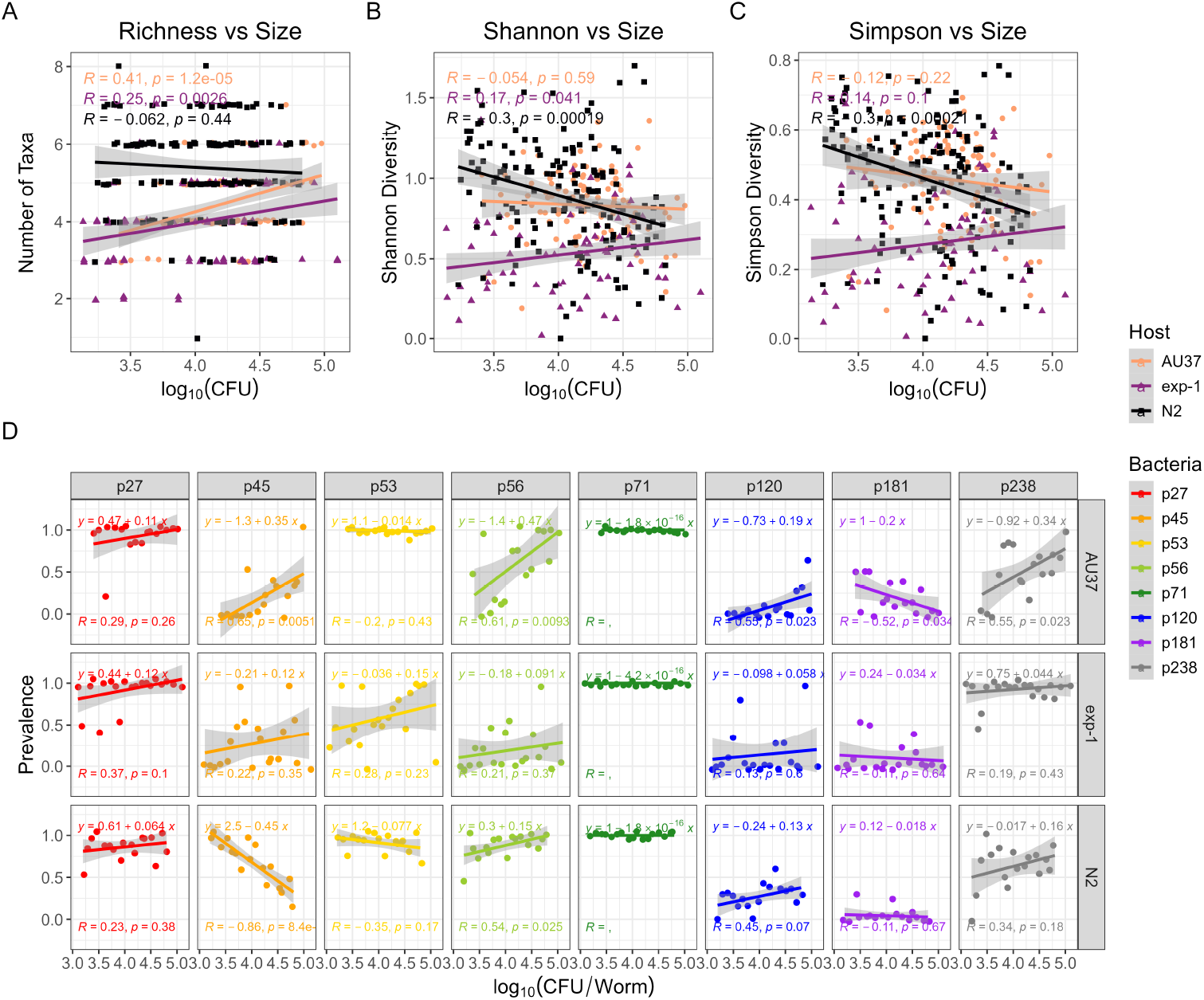
(A-C) Diversity vs. total bacterial load in worm-associated microbiota. Measures of diversity are shown in order of contribution of evenness to the metric: (A) Richness (number of taxa in individual worms); (B) Shannon diversity; (C) Simpson diversity. (D) Relationship between prevalence of individual taxa (fraction of individual hosts in which the taxon is present) and total bacterial load. Bacterial load was binned by rounding log-scale CFU/worm data to the first decimal place.

#### Model Expansions

##### Space Limited Growth without Interactions

To determine the role of interactions within the microbiome on the structure of the host-associated communities, we first developed a simple stochastic null model considering only migration, birth, death and competition for space within the host:

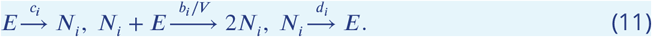

(This is also Eq. (1) in the *Main text*.) Here *E* represents an empty space and *N*_*i*_ a bacterium of type *i*. In the neutral model, all bacteria compete on equivalent terms; one bacterium of any type will occupy one empty space, and colonizing bacteria cannot be displaced through inter-species interactions. The first reaction corresponds to a worm host being colonized by a bacteria of type *i* and is characterized by the migration rate *c*_*i*_. The next reaction describes a bacteria of type *i* undergoing space-limited growth within the gut of its host with a maximum doubling rate *b*_*i*_. The last reaction corresponds to the death of the bacteria with a constant rate *d*_*i*_. The sum of all the empty spaces and bacteria within the gut is a constant equal to the carrying capacity *V*.

These reactions define a set of transition rates for the creation and destruction of a particular bacterial species i:

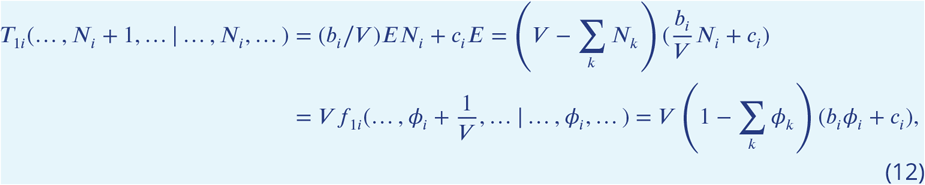

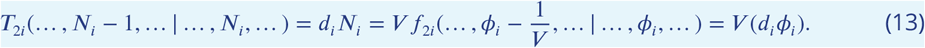

Here we have used the fact that the number of empty sites is equal to the carrying captivity minus the bacteria already existing in the gut. Additionally, we have expressed the transition rates in terms of population densities, *ϕ*_*i*_ = *N*_*i*_/*V*. From these reactions, we construct a Master Equation governing the time varying probability of having a certain number of bacteria within the gut of a *C. elegans* host:

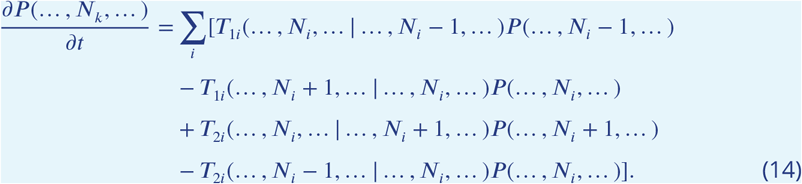

We perform the Kramers-Moyal expansion in the spirit of a Van Kampen system size expansion (***Van Kampen, 1992***) to recover differential equations governing the mean of the population densities of the bacteria populations and a Fokker Planck Equation governing the fluctuations about this mean. To do so, we rewrite the master equation in terms of densities *ϕ*_*i*_ using operator notation where 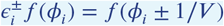.

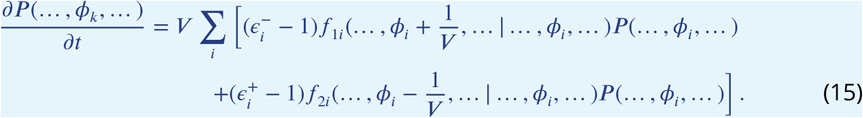

We expand the operator in a system size expansion with respect to the small parameter 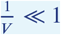

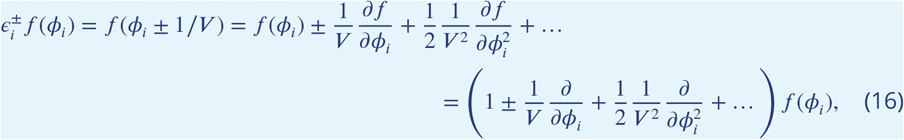

and

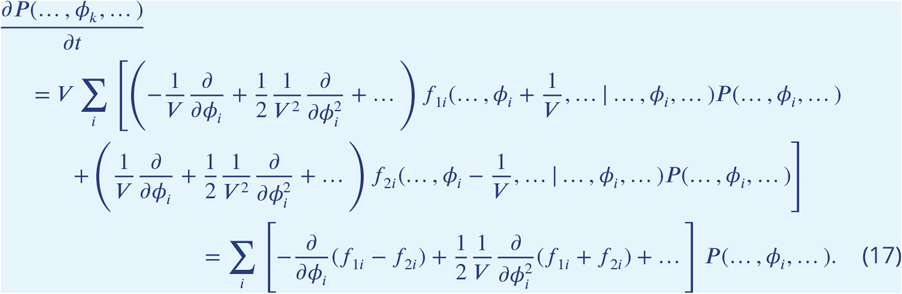

If we now truncate this expression to first order in the inverse system size, we obtain a Fokker Planck Equation. This Fokker Planck equation has a corresponding Langevin equation consisting of a mean field equation for the population density plus additive noise with a specific covariance structure. In general, for a Fokker Planck equation of the form 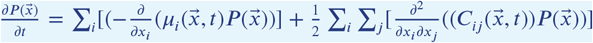, the corresponding Langevin equation is of the form: 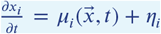, where *η*_*i*_ is additive noise with zero mean and covariance 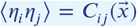. Therefore, the equations governing the means of the population densities *ϕ*_*i*_ is a generalization of the one used by ***Vega and Gore*** (***2017***) and is:

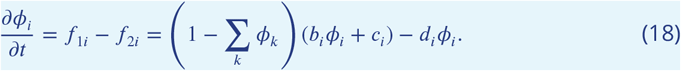

The predicted covariances for this simple model are

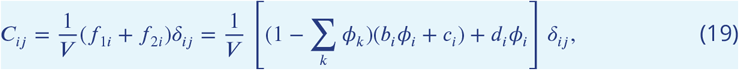

where *δ*_*ij*_ is the Kronecker delta function. This corresponds to an identity correlation matrix. This is also reproduced in the main text.

We can further validate our analytic calculations by performing stochastic simulations on this null model. We simulated the model using the Gillespie algorithm ***Gillespie*** (***1977***), which samples exact stochastic trajectories obeying the set of reactions we defined. To limit the total number of parameters in our fits, we chose to fix the birth rate and the migration rate for each bacterial species *i* to be the same. In Appendix Fig. 7, we show the calculated correlation matrix of multiple realizations of our stochastic simulation for the best fitting parameters. The simulation is able to capture the mean of the data. Additionally, it shows that, for this null model, there can be no off-diagonal correlations that are significantly nonzero. This validates our analytic calculation.

**Appendix 1—figure 7.**
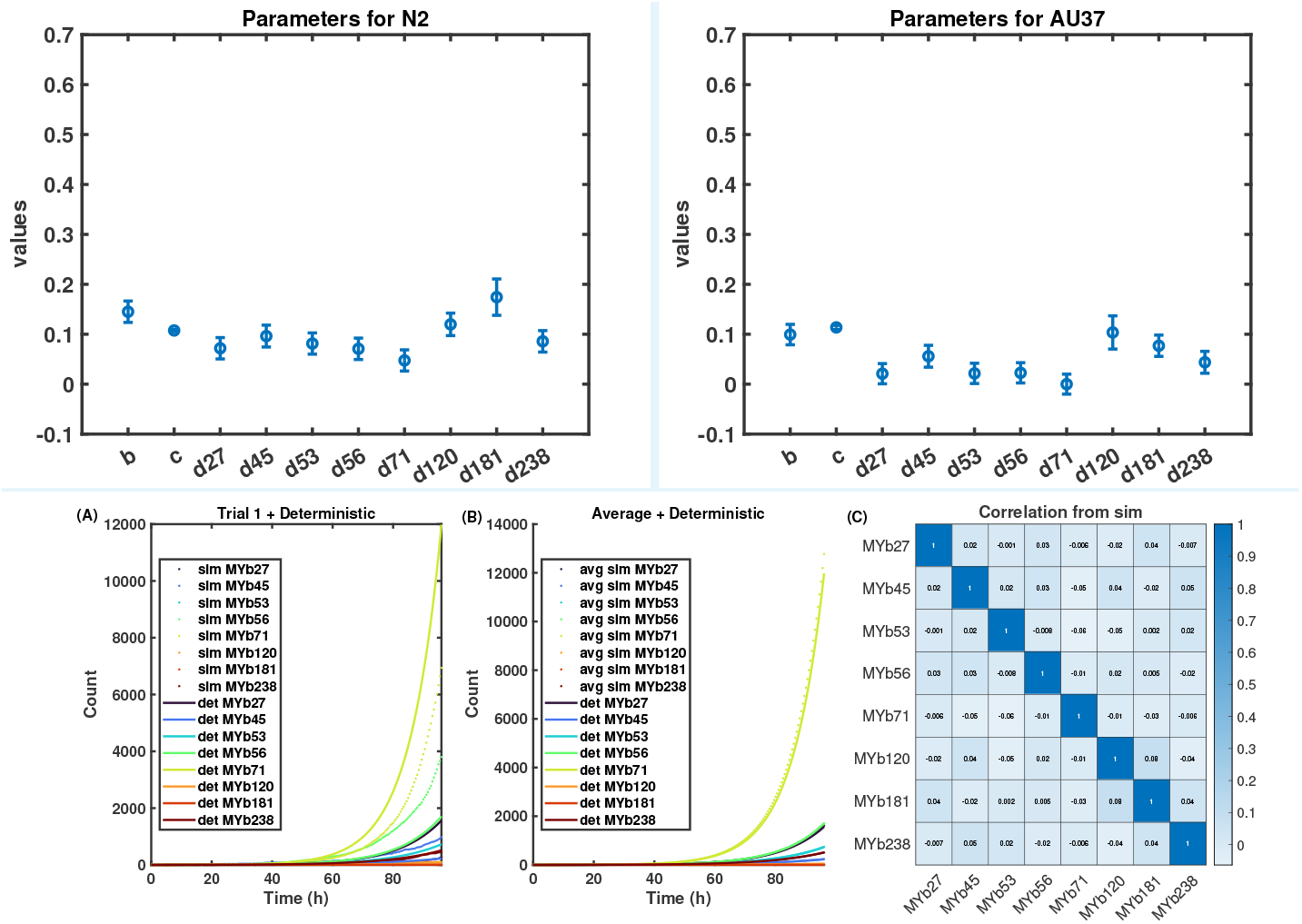
Top: Best fit parameters for N2 and AU37 worms. Bottom: Simulation of null model with parameters fitted to match the mean and variance of the bacteria species abundance within a worm N2. **(A)** Sample trajectories of bacteria concentrations for one trial of the stochastic simulation compared to the theoretical deterministic mean field. **(B)** The average trajectory of bacteria abundances of 1000 simulated trials compared to the mean field theory. **(C)**The calculated correlation matrix of the simulated trials.

##### Bacterial interactions

As an expansion of the simple model, we propose a model incorporating pairwise interspecies interactions. These interactions can be broadly classified into three categories, of which other plausible interactions represent special cases: warfare (decrease both species number by one), predation (increase one species while decreasing the other species), and facilitation (increase both species by one).

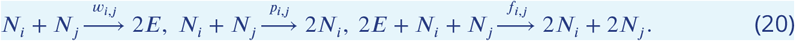

Using the same steps as outline in previous section, we calculated a master equation for the probability of abundances of the different bacteria, and performed the Kramers system size expansion to find a corresponding Fokker-Planck equation and the corresponding Langevin equation governing the dynamics of the abundances. The resulting equations governing the mean abundances are:

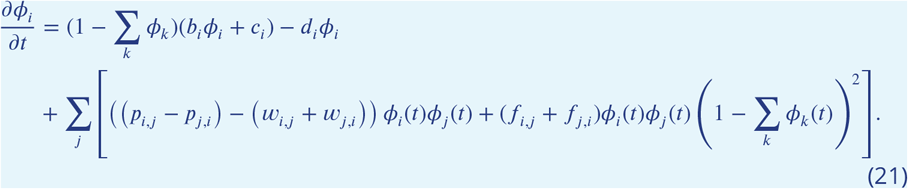

The resulting covariances (not correlations!) for *i* ≠ *j* within the model are:

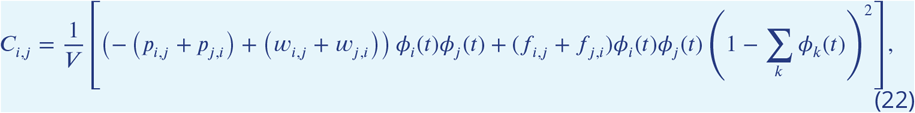

where *ϕ*_*i*_(*t*), again, are the ensemble-mean abundances of the species in the units of the carrying capacity, which solve the mean field equations for the model. For *i* = *j*, the diagonal covariances are:

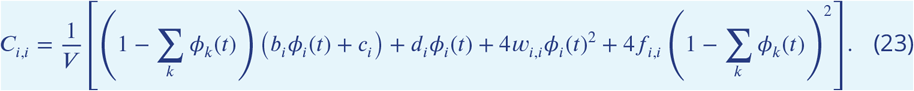

A prediction of this model is that the off-diagonal covariance scales as the product of the mean abundances *ϕ*_*i*_(*t*)*ϕ*_*j*_(*t*). In Appendix Fig. 8, we test this prediction by dividing each covariance term by the mean abundances of the contributing bacterial taxa. If there is no facilitation, our model predicts that the ratio of the off-diagonal covariance elements to their means should remain constant with time. Here we observe that this ratio on average decreases slightly with time, consistent with expectations for the contribution of facilitation to the covariance terms. Since the number of parameters are too many to fit for the available data, we can further simplify the model by letting the warfare, predation, and facilitation be the same values for each interaction *w*_*i*,*j*_ = *w, p*_*i*,*j*_ = *p, f*_*i*,*j*_ = *f*. Doing this and fitting the average off-diagonal covariance values we find 2 · (*w* − *p*) = 0.2, 2 · *f* = 6.7, and *V* = 2.1 · 10^4^. Which indicates that in this simplified model the warfare is greater than predation.

**Appendix 1—figure 8.**
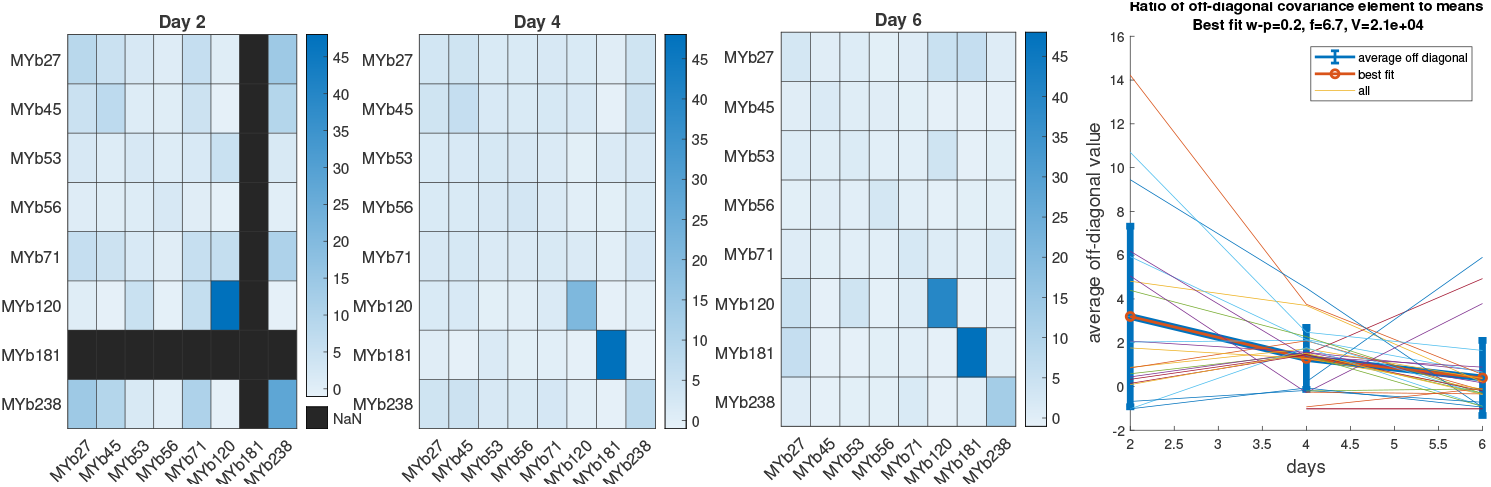
The ratio of *C*_*i*,*j*_ /(*X*_*i*_*X*_*j*_) for N2. Most of the off-diagonal elements are positive indicating that in the interacting model the warfare term and facilitation are greater than the predation terms. Ratio of off-diagonal elements to their corresponding mean abundances *C*_*i*_*j*/(*X*_*i*_*X*_*j*_).

##### Host Heterogeneity

Derivation of relationship between data covariance and parameter covariance matrices

Assuming that bacterial abundances are a deterministic function of the parameters used to initialize their growth and further assuming that the parameters are drawn from a Gaussian distribution centered at their best fit values, 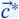 i.e. 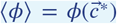, we can calculate a relationship between the covariances of the parameters and the abundance covariance matrix. Using a Taylor expansion of *ϕ* with respect to the parameters 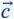 centered around 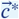 to the first order, 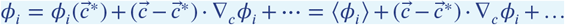, we can calculated the data covariance matrix. Let the matrix element *G*_*k*,*j*_ be defined as 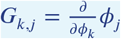. Then

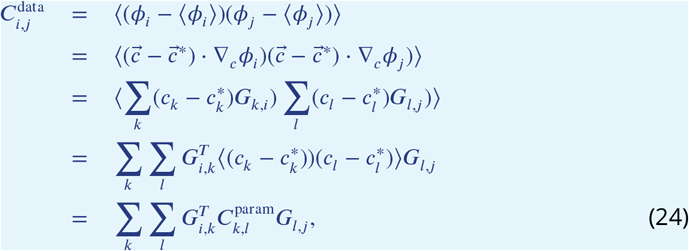

which is the result *C*_data_ = *G*^*T*^ *C*_param_*G* found in the *Main Text*. It is then a straightforward linear algebra problem to solve for *C*_param_ in terms of *C*_data_, as we did in the *Main Text*. However as *G* is not necessarily full rank, pseudo inverses must be used.

As mentioned in the *Main Text*, empirical calculation of the predicted parameter correlation matrix from data via the pseudo-inverse method requires the assumption that parameters are drawn from Gaussian distributions centered at their best-fit values. To test our Gaussian approximations for the distribution of parameters, we can find best parameter fits for the simple neutral model for bootstrapped resampled data. These fits tell us something about the uncertainty of the fitting parameters but should not be interpreted literally; instead, they serve as a rough description of the parameter space. The resulting histograms of best fit parameters are, in general, skewed and often multi-modal, contradicting our assumption of Gaussianity for for the distribution of parameters (cf Appendix Fig. 9). This either indicates that there are more complex density dependent interactions than we have modeled here, or that there are several sub-populations of distinct worms.

**Appendix 1—figure 9.**
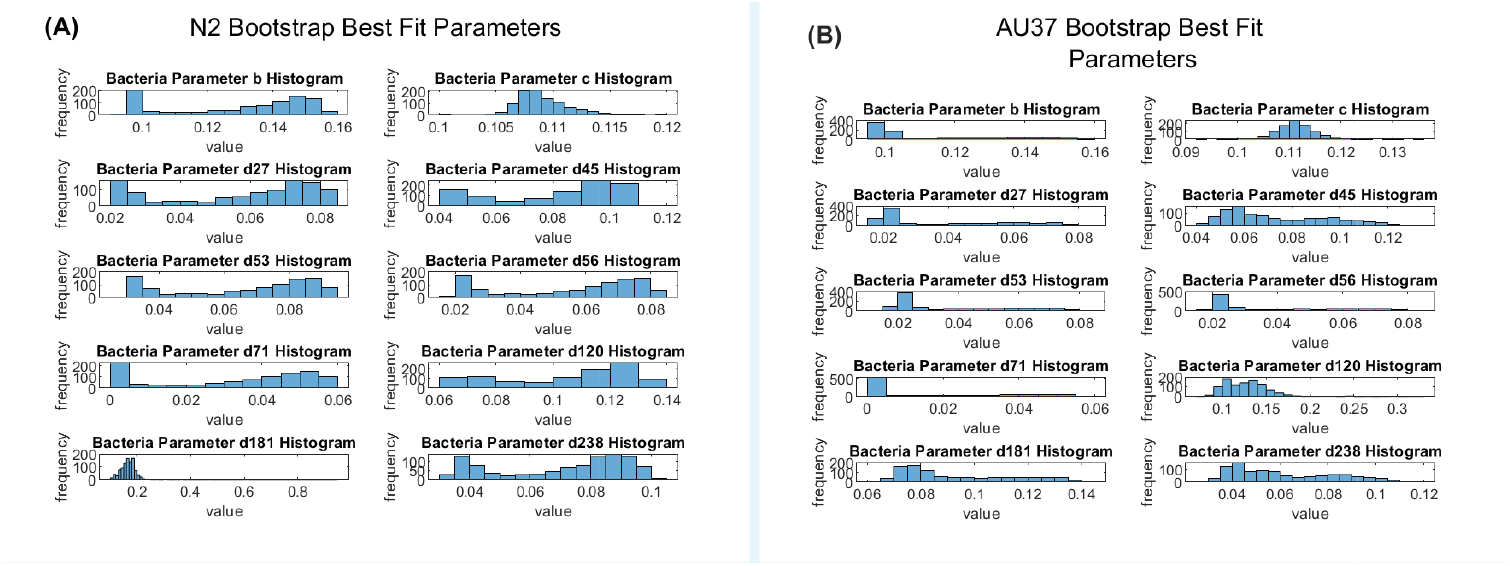
Distributions of parameter fits for (A) N2 and (B) AU37 microbiome communities.

**Appendix 1—figure 10.**
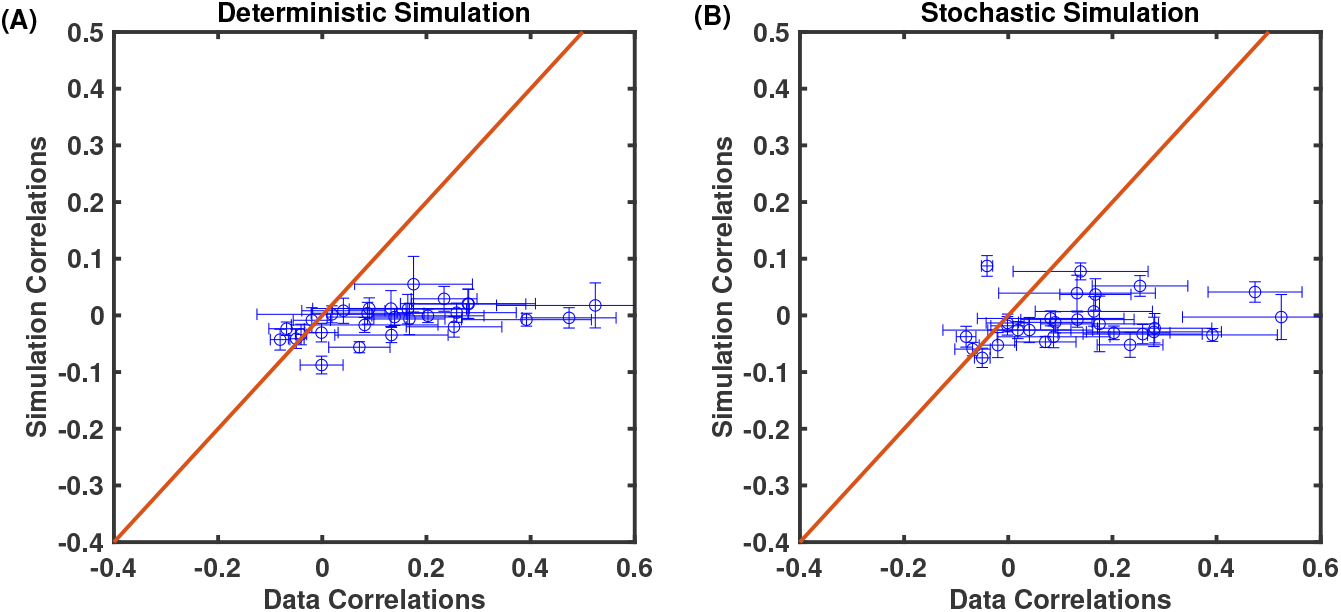
Off-diagonal correlations compared to data for simulations of both the deterministic and the stochastic model run with parameters drawn form uncorrelated Gaussian’s centered around best fit parameters.

